# Auditory-motor interactions are sensitive to adaptation and learning on multiple timescales

**DOI:** 10.64898/2026.03.23.712909

**Authors:** Haiqin Zhang, Giorgia Cantisani, Shihab Shamma

## Abstract

Auditory-motor learning is critical for mastering the production of complex sounds, such as speaking and playing music. It is anchored upon internal models of interactions between actions and their sensory consequences, which are fine-tuned by minimizing the errors between the predicted and received sound. Here, we investigated the neural dynamics of sensorimotor learning by manipulating sensorimotor surprisal, with high surprisal defined as unexpected changes in the mapping between actions and their auditory outcomes. Participants performed a piano-playing task in which the key-to-pitch mapping switched unpredictably among three configurations, creating a dynamic sensorimotor environment that required ongoing adaptation. At the change boundaries, a signature of violated motor-to-auditory predictions was found in the auditory evoked responses at N100, which could not be attributed to either purely auditory surprisals or motor execution errors. This prediction error is modulated by short-term context, with greater error responses following longer periods of no map change, indicating that the brain continuously tracks short-term map contexts and rapidly adapts to them. In contrast, 30 minutes of extended goal-directed training on a single key-pitch map modulated early ERP (0-60 ms) amplitude only for the trained map, a modulation that can be explained by a slow, training-driven update of the sensorimotor map. Hence, while auditory predictions from motor actions can be implicitly learned within short-term contexts for rapid adaptation, the complementary process of building a reliable sensorimotor map requires targeted training sustained over time. Our approach of studying auditory-motor surprisal in time-varying sequences reveals that auditory-motor learning is fast, context-sensitive, and shaped by both short- and long-term experience.

**Significance statement:** Understanding how the brain links motor actions with their sensory consequences is key to explaining how complex skills are acquired and subsequently adapted to changing environments. Yet, the neural mechanisms underpinning the evolution of internal sensorimotor associations across different timescales remain to be elucidated. Here, we extend the concept of *surprisal*, traditionally used in studies of perception, to the sensorimotor domain, where it is elicited by violations of expectations about the sensory outcome of an action. These expectations are generated by an internal model of sensorimotor associations that must nevertheless remain sufficiently flexible to enable rapid adaptation. Results show that surprisal responses are modulated differently by short-term sensory feedback and longer-term training, via two distinct neural mechanisms underlying adaptation and long-term sensorimotor map updates. These findings advance our understanding of the neural dynamics of sensorimotor learning and inform the design of emerging sensorimotor technologies, enabling novel forms of human-machine interaction and creative applications.

## Introduction

Continuous interactions between the sensory and motor cortical areas are fundamental for the execution of complex actions, such as speaking, playing music, writing, and balance control, among others. Control theorists described early how sensory feedback enables rapid and accurate adjustment of motor commands to achieve the intended outcomes and cope with noise (1; 2). In this context, action execution can be seen as a closed-loop iterative process within a sensorimotor system that combines contributions from both long-term knowledge accumulated through experience and short-term sensory feedback for fine control and rapid adaptation to a changing environment during performance (3).

Our work builds upon a framework referred to as the *Mirror Network* (4), that links control-theoretic principles (1; 2) to neural substrates for auditory-motor tasks, where the action-feedback loops reside in complementary pathways between motor and auditory regions in the cortex. This framework consists of two complementary pathways: (1) a *forward* pathway (decoder) from the motor to the auditory regions generating *predictions* of the sounds corresponding to the actions; and (2) an *inverse* pathway (encoder) from auditory to motor regions translating an intended sound into its corresponding motor commands (Figure 1). This internal model is *adaptive* and is learned by matching auditory predictions with sensory feedback (4). The resulting prediction errors yield a measure of movement accuracy that is then used to inform adjustments of the sensorimotor plan.

**Figure 1:**
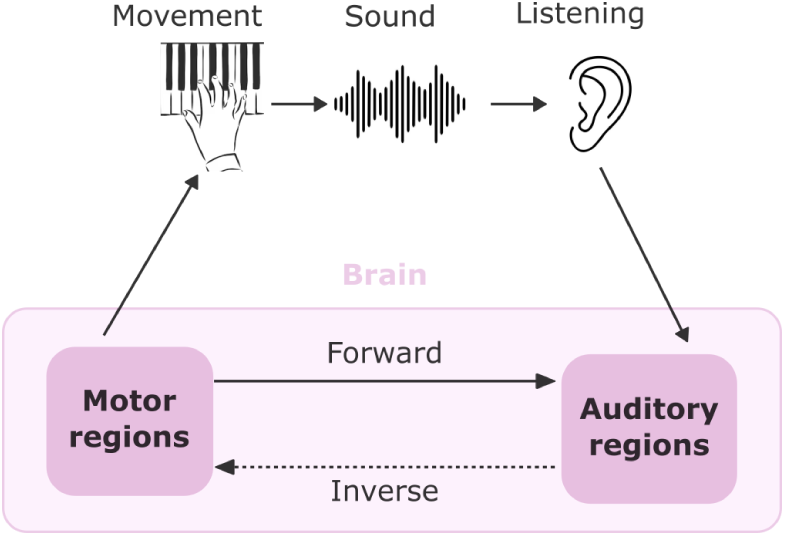
Schematic of the MirrorNet model proposed by Shamma et al., (4) in relation to the neural substrates of a sound production task. The *forward* pathway (decoder) maps the motor to the auditory regions, generating *predictions* of the sounds corresponding to the actions; the *inverse* pathway (encoder) maps auditory to motor regions, translating an intended sound into its corresponding motor commands. Predictions are informed by the previous context, *i.e.,* the previous actions in the motor sequence and the previous tokens in the auditory one.

A central question is how this framework can account for both the rapid adjustments required for adaptation to a changing environment and the stable maps acquired through long-term learning. In this work, we show that these two forms of learning–rapid adaptation and robust sensorimotor learning–arise from two distinct processes that update the sensorimotor map with different learning rates (raid vs slow learning) while integrating information at different timescales (short-term vs long-term context). As an example of rapid adaptation, compensatory speech responses to auditory feedback perturbations emerge within a single trial (5; 6; 7). Neural responses to such perturbations are observed in early ERP components, such as the N100 and other error-related negativities around 100–250 ms (8; 9). The timing of these error responses is consistent with neural signatures associated with the suppression of self-initiated sounds when there is a clear prediction of expected auditory feedback (10; 11; 12). On the other hand, mastering speech or a musical instrument requires years of training to build stable sensorimotor associations through repeated practice.

Taken together, these lines of thought suggest a separation between slow learning of stable sensorimotor maps and a fast, error-based correction system that adjusts existing maps in response to sensory feedback perturbations. Behavioral evidence supports the existence of these two processes: Van Vugt and colleagues showed, in an auditory-motor hand-reaching task, two dissociable mechanisms operating in parallel: one correcting movements based on feedback, and one encoding the movement required to reach a target in the first place (13). Similar dissociations have been observed in speech, where the timescale of adaptation depends on the nature and persistence of the perturbation. Some alterations of articulatory mechanics, such as those induced by a bite block, are compensated for almost immediately, whereas adaptation to others, such as artificial palates, develops only gradually over days or weeks (14). This difference suggests that the engagement of rapid compensatory mechanisms versus longer-term learning may depend on the severity and persistence of the perturbation.

Violations of such sensorimotor predictions^1^ have previously been probed with ERPs in classical oddball paradigms that introduce unexpected sensory feedback (15; 9; 8). These designs, however, did not control for the confound between auditory-motor and pure auditory surprisal driven by explicit target–response comparisons. Some experiments addressed this confound by comparing a simple key-pitch pairing that either stays constant, or changes randomly with each key press (8). However, each key press is still treated independently, precluding investigations of trial-to-trial effects related to long-term learning of a particular sensorimotor map and short-term adaptation to systemic perturbations – processes which are intrinsically context dependent.

Taken together, the existing approaches still provide limited insight into the continuous evolution of sensorimotor predictions. We addressed this gap by introducing a design in which sensorimotor predictions evolve continuously as an effect of both adaptation and training. This allows us to probe neural responses to violations of these predictions while dissociating the influence of short-term context from long-term learning in the following analyses:

### Experiment I. A variable map playing task to probe rapid adaptation

We introduce a continuous auditory-motor task—the *variable map playing* task—in which sensorimotor bindings are perturbed between periods of adaptation. Throughout the task, auditory-motor surprise is elicited by unpredictably changing the key-pitch mapping of a keyboard (Figure 2A) every few seconds. Between key-pitch mapping changes, the mapping remains stable for two to ten seconds (approximately corresponding to one to ten keystrokes). As we shall demonstrate, this design minimizes the contribution of purely auditory surprisal, ensuring that the observed electrophysiological response relates to surprisal elicited by the auditory-motor task.

**Figure 2:**
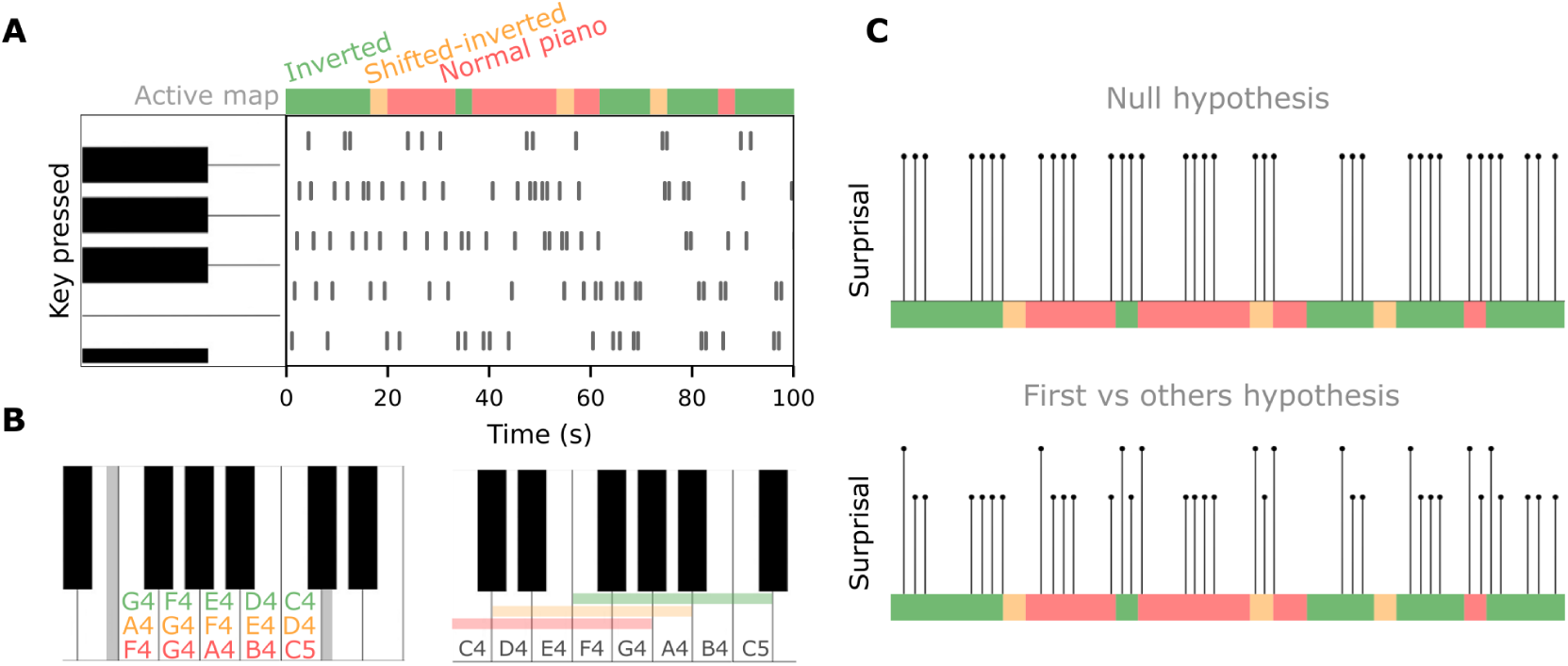
Experimental paradigm and hypotheses. **A.** Example of keys pressed by a participant over time. Vertical lines mark individual strokes, while the colored bar shows the active key-pitch map (green: inverted, yellow: shifted-inverted, red: normal). **B.** Key-pitch assignments on a standard piano keyboard, for each map. **C.** Hypotheses about surprisal differences between *first* and *other* keystrokes after a map change. The null hypothesis assumes no effect of the map change; the ‘first vs. others’ hypothesis predicts higher surprisal for the first keystroke than for later ones.

### Experiment II. Targeted task training to probe lasting sensorimotor map learning

Participants underwent a guided training session focused on learning to reproduce melodies on one constant key-pitch map (Figure 2C). After this training period, interactions between long-term training and short-term sensorimotor perturbations are examined in a second variable map playing session. We also considered the effect of baseline musical abilities, representing long-term sensorimotor knowledge, on the formation of the auditory-motor maps.

### Experiment III. Contribution of auditory and motor components to rapid adaptation and lasting sensorimotor learning

To assess the relative contribution of auditory and motor components to the neural signature of auditory-motor surprisal, we included an *auditory-only* task (passive listening) and a *motor-only* task (muted playing). Neural responses to these tasks were compared with those to the variable-map-playing task, which combines auditory and motor responses. We relate the auditory and motor components of the neural response to short-term adaptation and stable sensorimotor map formation, examining changes in these components before and after the target training in experiment II.

In our experiment, we combine a dynamically changing sensorimotor environment with the concept of *surprisal* to probe how internal motor-to-auditory predictions are updated across multiple timescales. Unlike traditional oddball paradigms, where deviance is typically defined by event probability within a fixed sequence, surprisal captures how unexpected an event is given its preceding context, even when the event itself is not inherently improbable.

From an information-theoretic perspective, surprisal can be modeled as *S*(*x | c*) = *−* log *p*(*x | c*), where a predictive model assigns probability to an event *x* given its context *c* (16). Statistical models of musical structure such as IDyOM (17) make predictions predictions by combining a long-term model trained on large musical corpora with a short-term model that adapts to the unfolding sequence, thereby capturing statistical structure across both long and short-term contexts. Surprisal measured by IDyOM has been used to show that neural responses to sounds are shaped jointly by immediate contextual dependencies and long-term exposure to transition statistics in both speech and music (18; 19).

Here, we extend the concept of surprisal to auditory-motor production. In this formulation, the context consists of previous keystrokes—both over short (recent actions) and long timescales (lifelong experience)— together with their associated auditory feedback, while the event corresponds to the present keystroke and its auditory feedback. If the auditory feedback matches what the expected sound, then the surprisal should be low. As with purely perceptual auditory surprisals, we expect auditory-motor surprisals to be driven by both short-term sensory feedback and by long-term knowledge acquired through sustained training. We further emphasize a key distinction between the well-established *auditory surprisal*, which reflects the violation of predictions formed from sensory context alone (18; 19) and the *auditory-motor surprisal* studied here, which reflects violations of predictions conditioned on both motor commands and the preceding motor–auditory sequence. As a consequence, auditory-motor surprisal is primarily relevant for self-generated actions and is not expected to arise in the same way for externally generated sounds.

In the absence of a model providing a continuous estimate of keystroke-level surprisal, we operationalize auditory–motor surprisal using experimentally defined proxies based on our variable-map playing design. Specifically, we treat the first notes following a key-pitch map change (referred to as *first* keystrokes) as higher-surprisal events relative to subsequent notes within the same key-pitch map (referred to as *other* keystrokes), which are expected to be more predictable once the current mapping is established. We then contrasted the neural responses to these two classes of keystrokes (Figure 2A). In addition, we compare neural responses to keystrokes that conform to a learned sensorimotor map against those arising from alternative key–pitch mappings, before and after sustained training. Our hypotheses were the following:

1. The *first* keystroke after a map change will violate the auditory prediction induced by the short-term context and therefore elicit a stronger surprisal response than *other* (following) keystrokes (Figure 2B). The surprisal response to upcoming events will decay rapidly as adaptation takes place.
2. Training on a specific key-pitch map will strengthen the motor-to-auditory predictions, such that keystrokes within this trained map will be *less* surprising than those played within untrained ones.
3. Since our task violates predictions by altering auditory feedback rather than introducing motor perturbations, neural markers of auditory-motor surprisals will be reflected in the auditory component of neural response.

## Results

### Experiment I

In the *variable map playing* task, participants were asked to freely improvise a series of melodies on a keyboard whose key-pitch map changed unpredictably every 2–10 seconds (Figure 2A). The playing session lasted 10 minutes without intervention from the experimenter. Participants were instructed to vary the melodies played, keep individual melodies short (approximately 4 notes), and avoid repeating the same notes or patterns. An example recording detailing the map changes, along with the auditory output, is available in the supplementary materials (Figure S2; supplementary video).

#### Neural signatures of auditory-motor surprisal

We began with the hypothesis that the evoked responses to the *first* keystrokes after a map change would elicit stronger auditory-motor surprisals than those of *others* due to the change in the key-pitch map (Figure 2A-B). To test this, we compared amplitudes of note-onset ERPs of the two classes of keystrokes (Figure 2C). To distinguish contributions of purely auditory versus auditory-motor surprisal to the evoked responses, we conducted a control experiment in which a set of naïve participants passively listened to the audio recordings of the notes played by participants in the main experiment (referred to as the *listening* condition, as opposed to the *playing* condition).

ERPs for for *firsts* vs *others* during playing are shown in Figure 3B. Statistical tests were computed on differences between *first* and *other* ERPs (Δ*_f−o_* difference waves) for both playing and passive listening. While Δ*_f−o_* was significant at 0.078-0.117 s for the playing participants (*i.e*., time points surrounding the N100 peak), listening participants only showed a significant Δ*_f−o_* at 0.218 - 0.273 s (Permutation cluster test, Figure 3B). In analyzing the scalp topography at N100, we found significant involvement of centrofrontal electrodes (Figure 3C; non-significant channels masked to white; *p <* 0.05, FDR-corrected Wilcoxon signed-rank test). These findings were consistent with the results of the spatiotemporal permutation cluster analyses of Δ*_f−o_* during playing and listening, which further revealed distinct scalp distributions for the difference waves around 100 ms during playing and 200 ms during listening (Fig. S3).

**Figure 3:**
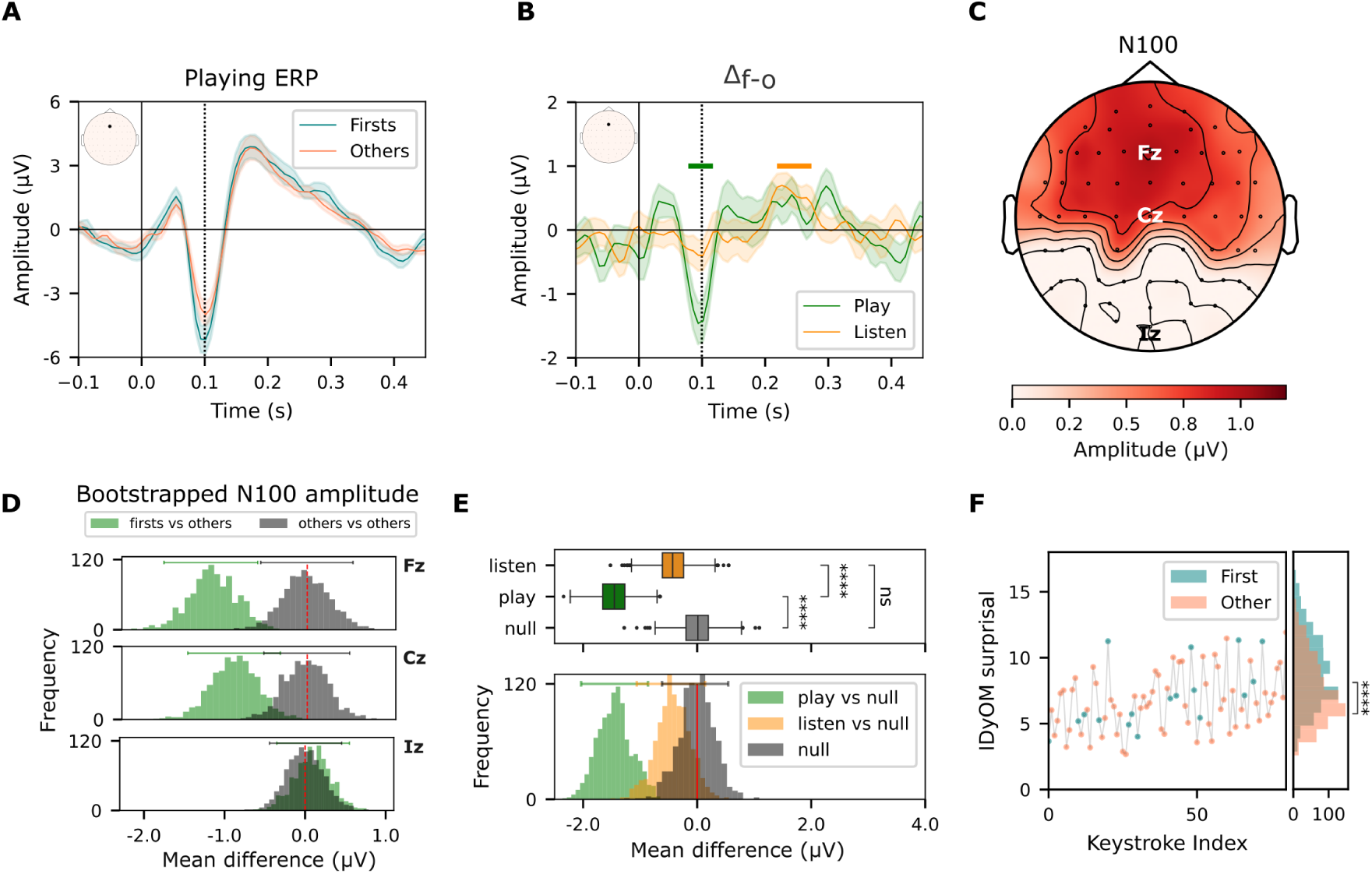
Neural signatures of auditory-motor surprisal. **A.** Grand-average ERPs of *firsts* and *others* at Fz. The dashed line marks 100 ms, where the N100 peak is found. **B.** Δ*_f−o_* difference waves. Bars indicate significant temporal clusters identified by the cluster-based permutation test (*p <* 0.05). **C.** Topography of N100 amplitude differences between *first* and *others* (Δ*_f−o_*), with non-significant channels masked to white (FDR-corrected Wilcoxon signed-rank test). **D.** Bootstrapped Δ*_f−o_* distribution (100-sample resamples, one from *firsts* and one from *others* over 1000 iterations) versus a null distribution where both samples are drawn from the pool of other keystrokes only. Horizontal bars indicate the CI95. **E.** Distribution of N100 Δ*_f−o_* in the playing and listening experiments, compared with the null distribution; bars mark significant clusters from the permutation cluster test (*p <* 0.05). **F.** Surprisal of heard notes as calculated by cross-validation using IDyOM: the scatter plot shows an excerpt from one recording, the histogram shows surprisal distributions over all recordings for all *firsts* and a size-matched random sample of *others* (*p <* 0.001).

Since there are more *other* keystrokes than *firsts*, we followed up with bootstrapping analyses to ensure that the Δ*_f−o_* is robust to unbalanced classes. This analysis consists of repeatedly sampling equal numbers of *first* and *other* keystrokes and computing the difference of means at each iteration. Bootstrapped difference distributions are shown for three representative channels relative to the region of significant electrodes: Fz within the region showing the most marked difference, Cz at the edge, and Iz outside showing no difference (Figure 3D). Figure 3E likewise compares bootstrapped distributions of N100 Δ*_f−o_* at the Fz electrode during playing and listening. Remarkably, only the playing distribution exhibited a significant difference from the null distribution (*p <* 0.001, empirical *p*-value). Playing and listening distributions were also significantly different (*p <* 0.001, permutation test, Figure 3E).

To investigate why the only listening condition had a cluster of interest at 200 ms, we examined the statistical properties of the note sequences themselves by evaluating the sequential surprisal of each note in the played melodies using IDyOM (*Information Dynamics of Music*), a statistical model of musical structure that causally estimates the probability distribution over all possible note-pitch continuations based on previous context in the sequence (20). Interestingly, a permutation test on IDyOM surprisals values revealed that notes produced by *first* keystrokes were on average more surprising than *others* (*t*-test, *p <* 0.001, Figure 3F), although the range of overall surprisal values did not differ.

#### Context-sensitive formation of auditory-motor associations

We next tested whether Δ*_f−o_* at N100 was modulated by the number of keystrokes played in the map preceding the map change. Thus, we defined a *0-keystroke* context as the case where the note immediately preceding the current *first* keystroke was itself a *first* keystroke in a previous map. A *1-keystroke* context corresponds to a case in which only one non-*first* note precedes the *first* keystroke and a *first* keystroke occurs two keystrokes prior, and so on. We hypothesized that *first* keystrokes with a longer context in the previous map would be more surprising and thus exhibit a larger N100 relative to *others* (Figure 4A). We fit a linear mixed-effects model to determine the relationship between keystroke context and Δ*_f−o_* at N100 for *first* keystrokes with different context lengths. We found that the amplitude of first-keystroke responses (*z* = *−*2.49, *p* = 0.014) decreased systematically with the number of preceding keystrokes in the previous map, and that the intercept of the model is significant (*z* = *−*2.46, *p* = 0.013), indicating a significant difference of N100 between all *firsts*, regardless of context, and *others*. Furthermore, consistent with the significant intercept, we found that when independently analyzing Δ*_f−o_* at N100 between each context category of *firsts* and the average amplitude of all *others*, the difference was significant for each context category (*t*-test, *p* = 0.005 for *0-keystroke*, *p <* 0.001 for *1/2/3-keystroke*, Figure 4B). Thus, *firsts* are significantly more surprising than *others* regardless of the number of keystrokes in the previous map.

**Figure 4:**
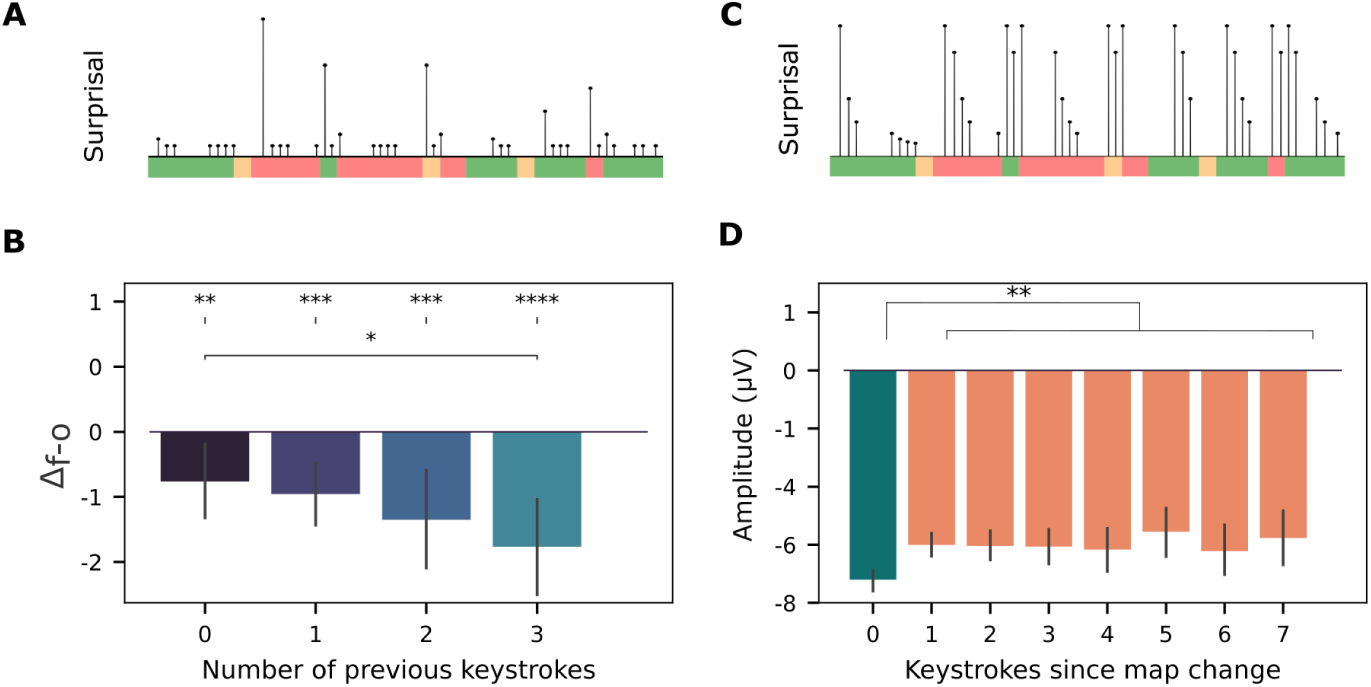
The influence of short-term context. **A.** Hypothesis: surprisal is modulated by the number of keystrokes in the preceding map. **B.** Bar plot showing mean difference in N100 amplitude between *first* and *other* keystrokes (Δ*_f−o_*) as a function of the number of keystrokes in the previous map. Independent samples *t*-test for comparisons between *firsts* with different numbers of previous keystrokes (bottom stars); one-sample *t*-test for comparisons between *firsts* and others (top stars). **C.** Hypothesis: surprisal is modulated by the number of keystrokes since the *first* keystroke in the current map. **D.** Δ*_f−o_* sorted by the number of keystrokes since the map change. Independent samples *t*-test.

In a complementary analysis, we examined the surprisal of *other* keystrokes as a function of the number of keystrokes since the last map change, with the hypothesis that it would decrease with a certain decay factor (Figure 4C). We categorized each keystroke based on the number of preceding keystrokes since the last map change (*firsts*, by definition, have no preceding keystrokes), and again fit a linear mixed-effects model to determine the relationship between number of preceding keystrokes and N100 amplitude of *other* keystrokes. We found no effect of the number of keystrokes since map change on N100 amplitude (*z* = *−*0.833, *p* = 0.405, Figure 4D).

### Experiment II

#### Effects of targeted skill-acquisition training

A 30-minute training session with a fixed key-pitch map (the *inverted* map) was included to assess the effects of extended skill acquisition on surprisal responses. The training consisted of imitation trials in which participants had one attempt to play 4-note melodies immediately after hearing them once (Figure 5A). Imitation trials were divided into two identical blocks with increasing difficulty within each block. To measure learning and engagement, each melody was scored based on how accurately it matched the target one (Figure 5B). Scores were significantly higher in the second than in the first block of training (Wilcoxon signed-rank test: *W* = 32.0, *p* = .002; Figure 5C).

**Figure 5:**
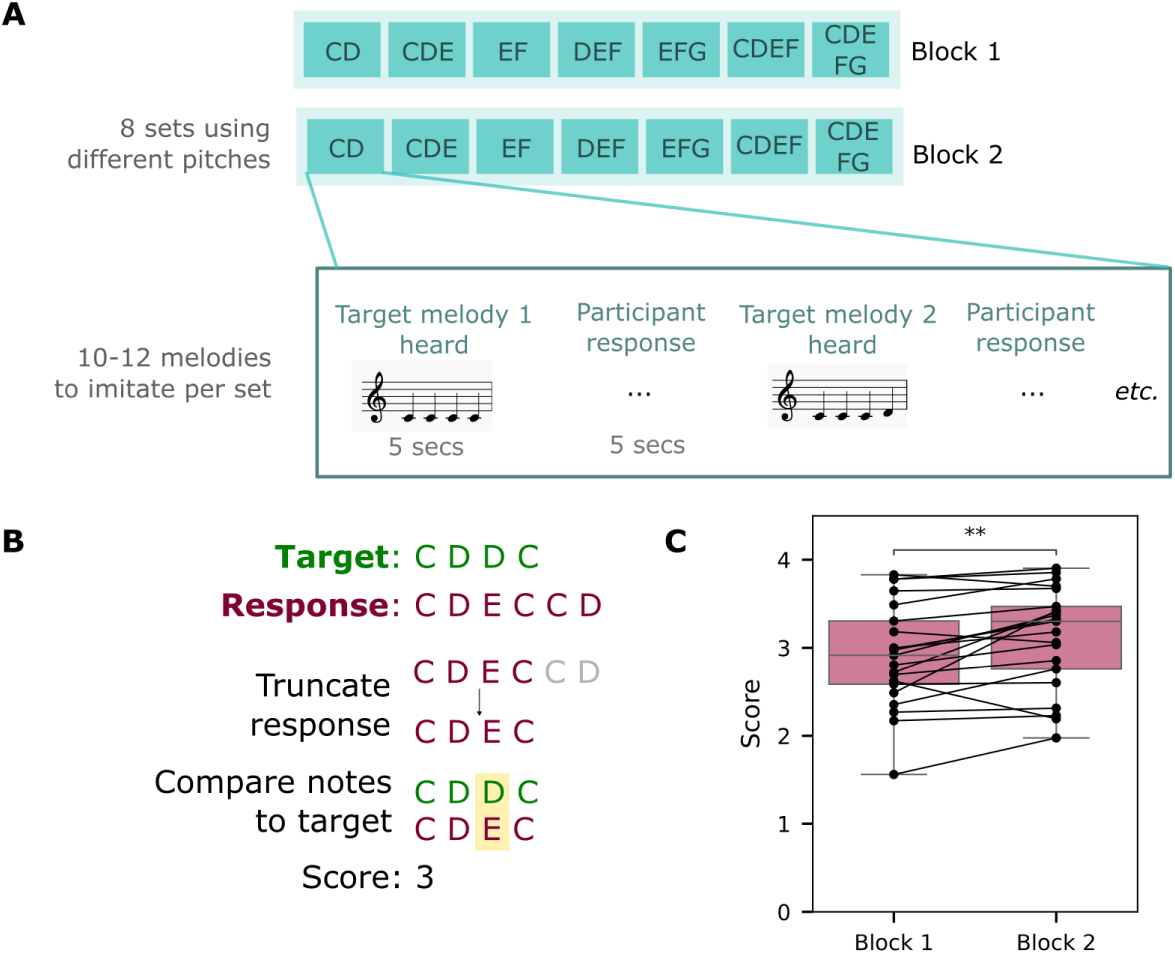
**A.** During training, participants imitated 4-note melodies by playing them back on the keyboard, in two blocks of increasing difficulty. **B.** Training was evaluated note-by-note against the target melody, ignoring rhythm and timing. **C.** Mean imitation scores for each block show learning over time (Wilcoxon signed-rank test, *p* = 0.002.). Lines denote individual participants.

The impact of extended training on auditory-motor surprisals was determined by comparing the *first* and *others* ERPs (Figure 6A, B). While a significant temporal cluster in the differences was found at 0.0781-0.117 s in the pre-training, a larger cluster was found post-training, from 0.047–0.109 s. Difference waves (Δ*_f−o_*) for each map were also plotted separately in pre- and post-training. No significant clusters were found in the pre-training in any map (note that there are smaller sample sizes since the ERPs have been subdivided into different maps). Post-training, Δ*_f−o_* in the normal map was significant at 0.070–0.109 s (Figure 6B-C). We then examined the difference in (Δ*_f−o_*) from pre- to post training. When sorted by the key-pitch map that was being entered into after a map change (Figure 6D), we found a significant difference at 0.007-0.054 s only for the *inverted* map. Topomaps revealed significant Δ*_f−o_* in central-frontal electrodes in both pre- and post-training for the N100, as well as the in the post-pre (Δ*_f−o_*) difference at 0-60 ms in the *inverted* map (FDR-corrected Wilcoxon signed-rank test, masked to show significant electrodes, significance threshold at *p* = 0.05, Figure 6E).

**Figure 6:**
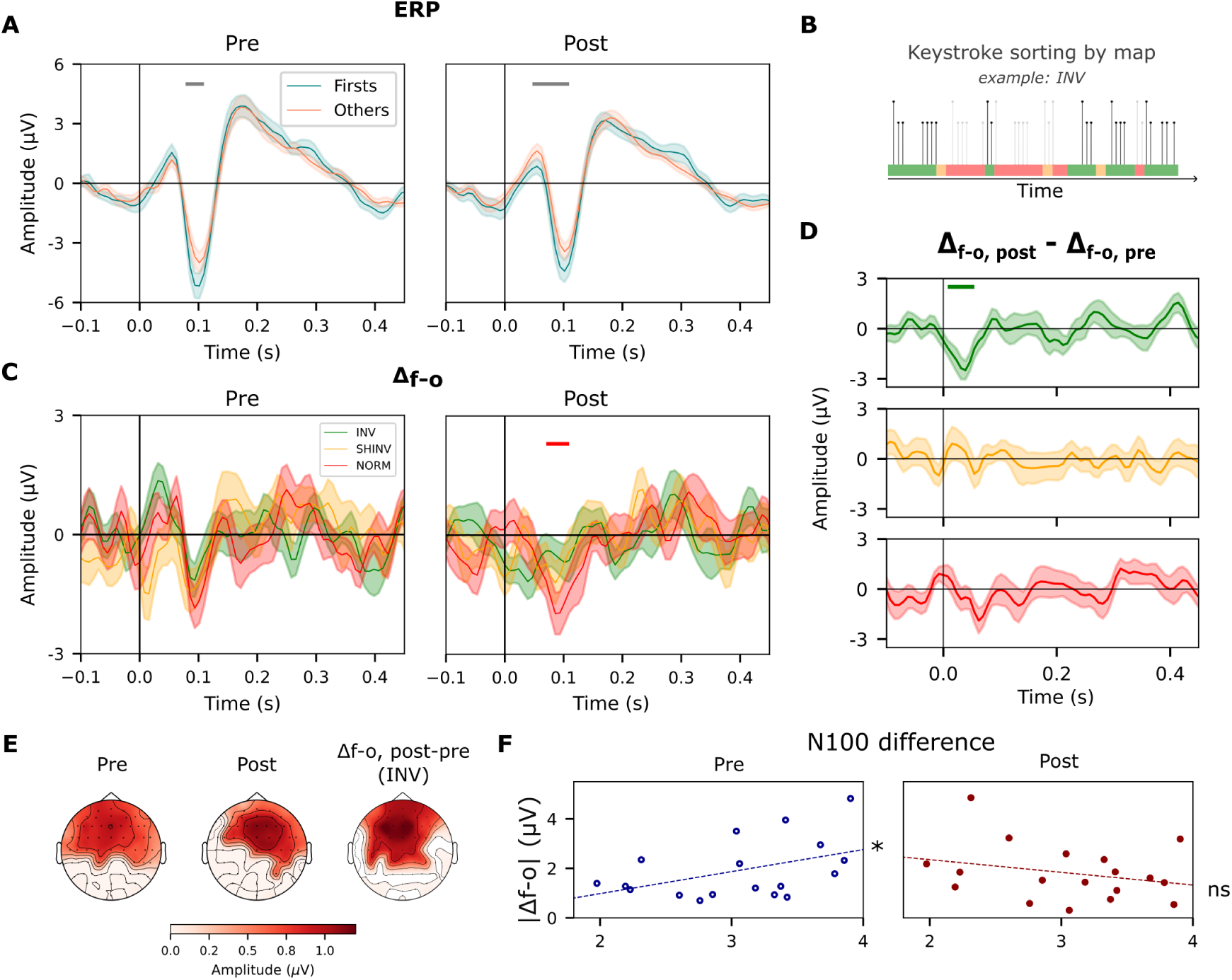
Effects of targeted and sustained training. **A.** Grand-average ERPs of *firsts* and *others* at Fz before and after training. Mean with shading representing SEM. **B.** Illustration of sorting keystrokes by the map active at the time of the keystroke. Colours represent active maps over time. **C.** Δ*_f−o_* in pre- and post-training at Fz, separated by the map at the time of the keystroke. **D.** Difference of Δ*_f−o_* waves in pre and post training for keystrokes separated by map. Bar represents significant time windows *p <* 0.05, 1-sample cluster permutation test. **E.** Topomap of Δ*_f−o_* at N100 in pre- and post-training, as well as the post-pre difference (Δ*_f−o_*, _post_ *−* Δ*_f−o_*, _pre_) for keystrokes in the trained map at 0-60 ms. Values represent averages over significant time windows. Colors are masked to show channels with significant Δ*_f−o_* (FDR-corrected Wilcoxon signed-rank test, non-significant channels masked to white, significance threshold at *p* = 0.05). **F.** Correlation between training score and Δ*_f−o_* at N100 in pre-training (*R* = 0.477*, p* = 0.0451, Pearson’s correlation) and post-training (not significant). Points represent individual participants.

#### Effects of musical expertise

Lifelong musical experience, including formal musical training as well as more general musical sophistication of non-musicians, can be considered as another form of training that preceded our recording sessions. We hypothesized that these factors may enhance participants’ ability to build auditory-motor associations and, therefore, their sensitivity to auditory-motor map violations. We use the training score of our learning task as a proxy for musical training, assuming that a higher musical background leads to higher scores in our learning task. Specifically, we measured the correlation between Δ*_f−o_* at N100 and the musical ability as measured by the training score in the pre-training period, and found a significant relationship (Figure 2F, Figure 6F; Pearson’s *r*, *r*(17) = 0.48*, p* = 0.045). The same analysis during the 0-60 ms period, split by the active key-pitch map, did not yield any significant correlations (Figure S7).

Finally, we performed the same correlation analysis using subject scores on the Goldsmiths Musical Sophistication Index. Neither the overall score nor the sub-scores for perceptual abilities produced a significant correlation with Δ*_f−o_* at N100 (Figure S8).

### Experiment III

#### Disentangling auditory and motor components in the playing responses

We next sought to dissociate auditory and motor contributions to neural responses. To this end, we analyzed neural responses as participants completed additional passive listening and mute playing tasks (Figure 2C).

We first show that during ordinary keyboard playing, auditory and motor components of the neural responses are superimposed. This is reflected in the evoked responses of the three different conditions: *playing*, which generates both components; *listening*, which produces pure auditory responses; and *mute playing* evoking pure motor responses (Figure 7A). The playing ERP, therefore, can be viewed as a combination of these auditory and motor components. The simplest such combination is the linear weighting of the pure components, which optimally fits the composite of the playing ERPs as illustrated for the Fz channel in Figure S11. Responses of all EEG channels, and comparisons of responses pre- and post-training can be found in Figures S4 and S5).

**Figure 7:**
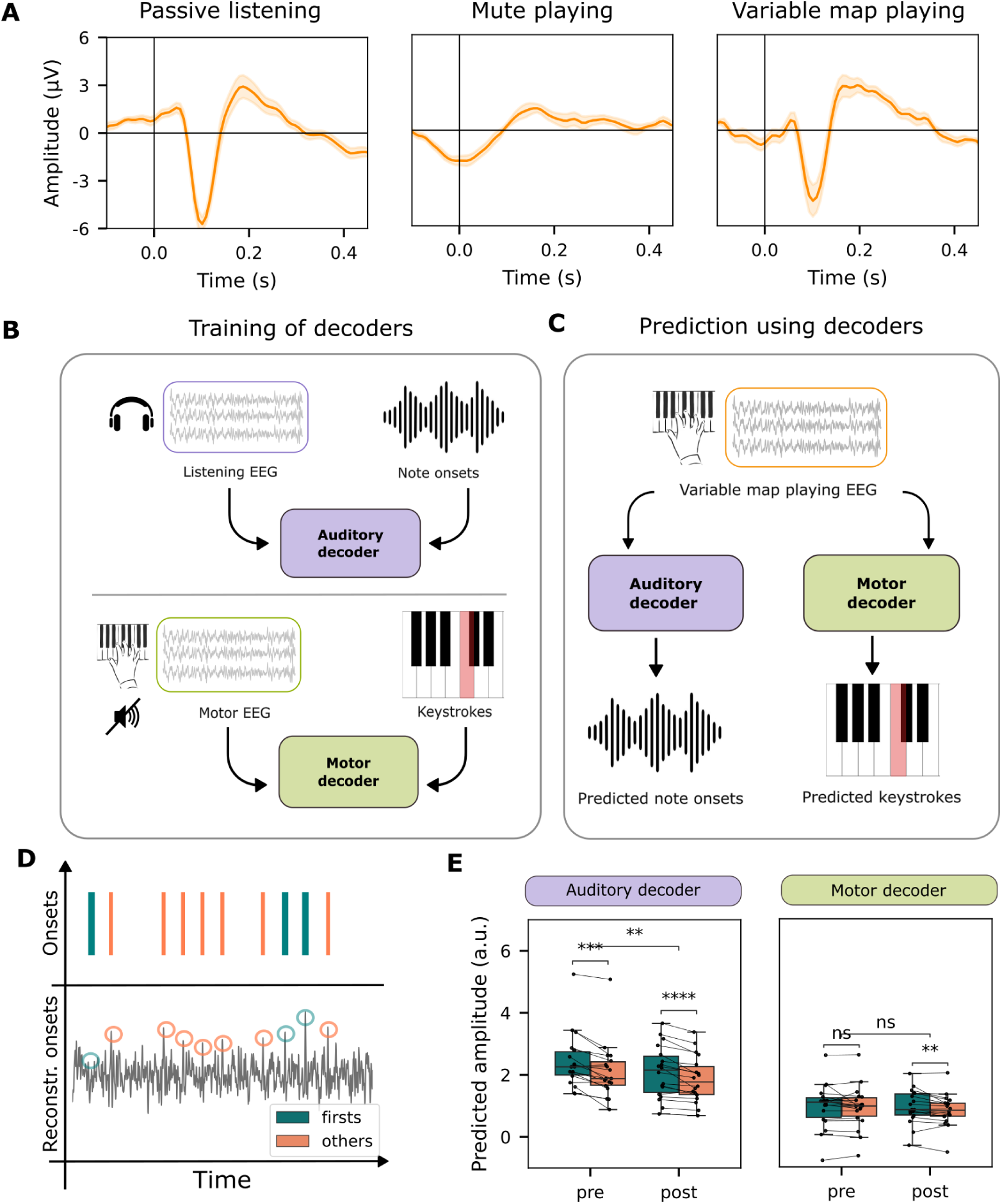
Disentangling auditory and motor components of playing responses. **A.** Grand average ERPs to note onsets at Fz for passive listening, mute playing, and variable map playing. **B.** The auditory decoder reconstructs note onsets from passive listening EEG; the motor decoder reconstructs key presses from mute playing EEG. **C.** Both decoders are then applied to the playing EEG. **D.** Example of ground truth and reconstructed onsets. **E.** Reconstruction amplitudes at *first* and *other* keystrokes are averaged and compared in pre- and post-training. Lines represent participants; bars show medians and quartiles. Wilcoxon signed-rank test, *p <* 0.001.

We used a decoding approach to disentangle the auditory and motor components in the playing (auditory-motor) condition and assess how they are affected by training. For each participant and condition (*i.e.,* auditory and motor), we learned the optimal mapping from time-lagged EEG activity to stimulus-related features via regularized linear regression (21) (Figure 7A). The *auditory decoder* was trained to reconstruct *sound onsets* from the passive listening EEG, and the *motor decoder* to reconstruct *keystroke onsets* from *mute* playing EEG (Figure 7B). Both decoders were then applied to *variable map playing* EEG (Figure 7C-D), and the mean amplitude of their predictions at note onsets was compared in pre- and post-training (Figure 7E). Note that the decoders are temporally agnostic, being trained on a wide time window (*±*500 ms around onsets), so they could not rely on EEG amplitude at a single latency.

Both decoders performed above chance in reconstructing note onsets and keystrokes, with higher amplitudes at note-onset times (Figure S9C-D). The *auditory decoder* predicted reliably greater amplitudes at *first* than at *other* keystrokes (main effect of keystroke type: *F* (1, 17) = 34.26, *p < .*001, Greenhouse-Geisser corrected), with no main effect of training or interaction. The *motor decoder* also predicted greater amplitudes at *first* keystrokes (*F* (1, 17) = 6.53, *p* = .020, Greenhouse-Geisser corrected), again without a main effect of training nor interactions. Yet, pairwise Wilcoxon tests on motor decoder predictions revealed that amplitudes at *first* keystrokes were greater than those of *others* in post-, (*p* = 0.00769) but not pre-training (Figure 7E, right panel), a result that will be discussed further in the next section. The same analysis performed on a size-matched samples of 100 firsts and others yielded the same conclusion (Figure S10). Overall, these findings suggest that auditory-motor prediction violations in the playing condition are mainly reflected in the auditory, rather than motor component of the neural response in pre-training, but both auditory and motor components contribute post-training.

## Discussion

Learning how to map movements onto complex sounds depends on a finely tuned communication between the motor and auditory systems. To probe how short-term sensory feedback and long-term knowledge each contribute to this cross-talk, we developed the *variable map playing* paradigm. This approach adapts the idea of auditory surprisal, well studied in passive listening contexts (18; 9), and applies it to self-generated sounds in which surprisal arises from violations of learned auditory-motor associations. Taken together, our findings show that auditory-motor predictions are modulated by both short-term sensory feedback and knowledge gradually accumulated through long-term training.

### The neural signature of auditory-motor surprisals

#### Auditory-motor surprisal is reflected in the modulations of the N100 component of the auditory response

One central hypothesis of our study was that keystrokes following a change in a key-pitch map elicit a stronger surprisal-related neural response than other keystrokes. We hypothesized that this surprise response is auditory-motor in nature (*i.e.,* due to a violation of the expected association between the motor action and resulting sound), rather than purely auditory (based on learned statistical patterns of note transitions).

Consistent with previous studies, we found a neural signature of auditory-motor surprise in the form of a significantly larger N100 in the ERP of *first* notes after a map change compared to that of *others*. The N100 is known as a component of auditory responses to sound onsets (22; 23; 24; 25), which is modulated during passive listening by contextual surprise within melodic sequences (referred to earlier as ‘pure auditory’ surprisals) (26; 18), as well as in speech (27). The modulation is explained by a model in which internal predictions of an incoming sound evoke a neural response of opposite polarity to that of encoding the sound itself. When internal predictions match the incoming sensory input, the two components partially cancel out, resulting in an attenuated net neural response. When they do not match, the N100 amplitude is enhanced proportionally to the mismatch or surprisal (18). This process also occurs with self-generated sounds, such as speech (28) or self-triggered tones (29; 10; 30; 11; 12), where N100 attenuation arises from accurate motor predictions rather than prior auditory context.

#### Effects at N100 cannot be explained by purely auditory surprisal

To distinguish between purely auditory and auditory-motor surprisal, we conducted a control experiment in which a separate group of participants listened to recordings of the original group’s playing sessions. The hypothesis was that such sequences should not exhibit a structural bias between *firsts* and *others*, and therefore should not elicit any contextual surprisal modulations when only heard. If the played melodies had exhibited meaningful structure, this could have biased the results, with the first note after a map change being, on average, more surprising than the others, and hence resulting in a confounding factor. Consistent with our hypothesis, we did not find any significant difference in the N100 between *firsts* and *others* in the listening (control) group, whereas such a difference was present in the playing group (Figure 3A-E).

Interestingly, however, we found a *firsts* vs. *others* difference in the P200 component among the control group, consistent with prior work associating this component with auditory surprisal (18). This difference was absent during variable map playing (Figure 3B), potentially because concurrent motor activity masked or modulated the auditory response. Supporting this interpretation, we found that both auditory and motor ERPs, obtained respectively from note onsets in the passive pure listening condition and keystrokes in the mute playing condition, exhibited clear P200 components (Figure 7A;S4), and that the playing ERP could be expressed as a linear combination of these auditory and motor components (Figure S11).

A computational analysis using IDyOM, a statistical model of musical surprisal (20; 31), converged on the same finding, yielding higher surprisals for *firsts* than for *others* (Figure 3F). We chose to model surprisal using IDyOM because its surprisal predictions have been shown to closely align with behavioral and neural responses (18; 17; 32; 33; 34). Several factors can at least partially explain such contextual surprisals, including interval leaps (the largest leap possible within the same map is only 7 semitones, whereas it may reach 12 semitones at a map change; Figure 2A), and musically surprising transitions (since participants were improvising the melodies they played, unexpected map changes may have rendered the melodies less musically coherent). Together, results indicate a small contribution of purely auditory surprisal to the neural response despite our precautions to limit it. This surprisal is nevertheless insufficient to account for the stronger N100 modulation in *firsts* observed during active playing, as shown by the comparisons with the listening control group (Figure 3B, E).

#### The surprisal effect cannot be explained by motor execution errors

In the MirrorNetwork framework (Figure 1 (4), the auditory prediction generated by the motor system is projected into auditory areas through a *forward* pathway (decoder), where it is compared with the actual sensory feedback. Prediction violations elicit a surprisal signal that is projected back to motor areas through an *inverse* pathway (encoder), supporting both learning and adaptation. In this model, auditory predictions can be violated in two ways: either the motor plant misses the articulatory target (*i.e.,* a motor error), or a change in the environment alters the sound outcome (*i.e.,* an auditory prediction error). Our experiment was explicitly designed to violate auditory predictions in a sensory-motor task by inducing an erroneous *auditory feedback* while maintaining a simple motor task that minimizes motor errors (single, slow finger movements).

To determine whether the surprisal signature could be attributed to an auditory or a motor process, we trained linear decoders to reconstruct note onsets from responses to passive listening and mute playing, and evaluated their transfer performance on playing data. We found that the decoder trained on passive-listening EEG data (purely sensory) could discriminate between *first* and *other* keystrokes in the variable-map playing data both before and after training. By contrast, a decoder trained with mute playing data (purely motor) could not detect a difference before training (the post-training results shall be discussed below) (Figure 7). We interpret this as follows: within the MirrorNetwork framework, the *forward* pathway is rapidly adjusted during initial exploration, generating accurate auditory predictions from motor commands before motor skills are refined. Thus, early surprisal responses reflect prediction errors in the auditory domain rather than motor execution errors.

### Sensorimotor interactions combine short-term sensory feedback with long-term knowledge

#### N100 reveals a continuous tracking of short-term sensory context for rapid adaptation

We extend previous findings showing N100 as a signature of auditory-motor surprisal by demonstrating that the brain progressively refines its auditory-motor model using information from each action. We hypothesized that short-term context primarily influences the early phase of sensorimotor learning or adaptation, as reflected in surprisal-related neural responses. During this phase, predictions are refined as movements are associated with corresponding sounds, causing surprisals to decrease steadily from one keystroke to the next. Once the auditory-motor map is acquired, any subsequent change in the map induces a new high-surprisal response that rapidly plateaus as auditory-motor predictions align with the new key-pitch map. Our analyses revealed that *first* keystrokes after a map change elicited significantly larger N100 amplitudes than subsequent ones (Figure3). Contrary to our hypothesis, however, the N100 amplitude of subsequent keystrokes remains relatively stable (Figure 4D). This suggests that after the surprisal response to the *first* note, the brain rapidly establishes accurate predictions for all other keys—either by extrapolating from the initial violation or by activating one of multiple stored auditory-motor maps. This interpretation aligns with findings from auditory feedback perturbation studies (35) in bilingual language production, where bilingual participants learn language-specific motor adjustments and apply them as a function of the language being spoken (36), as well as in audio-visual tasks with intermittent visual field perturbations (37; 38).

Interestingly, we found that *first* keystrokes elicited a stronger N100 response as the number of keystrokes played in a previous map increased (Figure 4B). We attribute this effect to the perceived stability of the short-term sensory context, *i.e.,* the more keystrokes occur without a map change, the more stable the context. Hence, a mapping violation is more surprising in a stable context than in one characterized by frequent changes. Prior work has theorized that the brain makes continuous predictions at multiple hierarchical levels. At a lower level, it predicts upcoming sensory events (e.g., individual note outcomes) based on the current sensorimotor mapping. At a higher level, it predicts when and how the sensorimotor mapping itself will change (39). Both prediction levels rely on sensory information, but operate on different timescales: immediate sensory feedback updates predictions about the next event, while accumulated sensory information over time updates the probability distribution governing potential map changes (40). This hierarchical predictive mechanism likely allows the brain to anticipate and prepare for changing consequences of motor actions.

#### ERP amplitude at 0-60ms is modulated by extended training distinct from N100 adaptation

Auditory-motor surprisal responses comprise distinct auditory and motor components that can be disentangled based on their differential evolution over 30 minutes of targeted training. As previously discussed, the *forward* pathway maps motor commands onto their predicted sensory outcomes, thereby generating auditory expectations. Violations of these expectations produce sensory prediction errors, *i.e.,* an auditory–motor surprisal. Such prediction errors drive rapid compensatory responses, which emerge even after only brief exposure to altered auditory feedback (5; 6; 7). In contrast, the *inverse* pathway maps desired sensory outcomes onto the motor commands required to produce them. Rather than generating sensory predictions, this pathway projects auditory prediction errors back to the motor areas to refine the sensorimotor mapping, enabling progressively more accurate predictions. Because learning this mapping requires solving a many-to-one inference problem (41), updates are expected to emerge gradually through repeated practice. Accordingly, we hypothesize that the later motor component of auditory–motor surprisal reflects the progressive updating of the sensorimotor map by projecting the error through inverse pathway, whereas the earlier auditory component primarily reflects violations of predictions generated by the forward pathway.

In our results, the N100 primarily indexes responses to short-term surprisal, whereas the amplitude from 0-60 ms is a marker of a lasting update to the sensorimotor map. Critically, these components exhibit distinct dynamics: the N100 shows rapid modulation during early exploratory exposure and returns to baseline after rapid adaptation, while the modulation at 0-60 ms only emerges following extended training. The interpretation of the N100 as indexing auditory surprisal along the *forward* pathway is consistent with our finding that, following training, a significant Δ*_f−o_* at N100 was observed only for keystrokes in the *normal* map, which was the most dissimilar to the trained map. This suggests that training on a markedly different map may have increased the auditory surprisal associated with the *normal* map.

In contrast, the 0-60 ms difference showed a significant modulation by training and was specific to the trained key-pitch map (the *inverted* map), indicating that this component of the neural response is modulated only after sustained exposure to the sensorimotor transformation. We thus interpret the modulation of amplitude at 0-60 ms as being related to arriving in a trained, familiar map rather than departing from a short term map still emerging after a few keystrokes of experience.

Together, these different ways in which amplitudes at 0-60 ms and N100 are modulated clarify the relationship between adaptation and learning: adaptation occurs within an existing predictive model supports immediate behavioral adjustment, but does not imply a change in the underlying sensorimotor mapping. By contrast, learning reflects the gradual acquisition of stable sensorimotor mappings. This distinction aligns with dual-process accounts of sensorimotor learning (13), in which a fast error-correction mechanism operates alongside a slower mapping system that supports robust, trained performance.

#### The effects of long-term musical abilities

We expected better musical abilities to enhance the participants’ capacity to build reliable auditory-motor associations and, therefore, their sensitivity to map violations. Musical training has been shown to improve abilities such as motor control (42; 43), auditory perception (44; 45), and audio-motor integration (46; 47; 48; 49; 50; 51; 52). These abilities possibly contribute to musicians’ ability to adapt faster to various environmental changes, such as rhythmic perturbations (53; 54; 55). Yet, the influence of long-term training on rapid adaptation in short-term, changing environments with perturbed auditory-motor associations remains unclear.

To further probe individual differences, we used melody imitation accuracy during training as a proxy for musical ability, as it is likely to be enhanced in trained or musically gifted individuals (56). We found that Δ*_f−o_* correlated with training performance, but only before training, suggesting that participants with higher initial skill may have stronger priors about the keyboard layout and were more adept at short-term auditory-motor adaptation, leading to larger prediction violations when the key-pitch mapping changed. Following training, this relationship disappeared, suggesting that any priors knowledge of keyboards was superseded rapid realignment of the internal model to the trained map.

We also examined associations between Δ*_f−o_* and musical sophistication as measured by the Goldsmiths Musical Sophistication Index (57). However, this questionnaire-based measure showed no consistent relationship with neural effects and was not retained as a primary index of musical ability. However, it should be noted that the questionnaire takes into account additional factors (e.g., emotional engagement and self-reported skills) that may be less directly tied to sensorimotor predictive accuracy than the training performance measured in our experiment.

#### Conclusions, limitations and future steps

Our findings reveal that auditory-motor learning operates over two distinct timescales. The brain rapidly and implicitly updates predictions of sensory consequences of actions within seconds, whereas refining motor commands from auditory feedback requires sustained, goal-directed practice over longer periods. This asymmetry between rapid adaptation and slow learning provides a mechanistic account of how complex sensorimotor skills are acquired and refined in domains such as music and speech.

In this work, we operationalized auditory–motor surprisal as a binary variable by contrasting expected and unexpected events induced by changes in the sensorimotor mapping. An important next step will be to develop computational models that estimate continuous auditory–motor surprisal from the full sensorimotor context, analogous to IDyOM for melodic surprisal (20) and probabilistic models of phonemic and lexical surprisal in speech (58; 59). Such models would allow prediction errors to be quantified on a trial-by-trial basis rather than through discrete experimental manipulations.

More broadly, future studies should move toward more naturalistic auditory–motor behaviors, where mappings between actions and sensory outcomes are high-dimensional, continuous and flexible, as in real music performance. This will require probabilistic models capable of learning the statistical structure of motor sequences and their associated sensory consequences, enabling continuous estimates of auditory–motor surprisal from ongoing behavior. Complementing these behavioral models, neural encoding approaches, such as temporal response functions (21), could be used to relate the estimated auditory–motor surprisal to continuous neural activity. Extending the framework further to incorporate motor variables—including finger, hand, and arm kinematics—would enable surprisal to be modeled jointly over auditory and motor domains, providing a fully bidirectional account of sensorimotor prediction and learning.

Overall, this works contributes toward a unified characterization of auditory-motor learning. Our results reveal distinct neural signatures of rapid prediction updating and slower sensorimotor map learning, providing evidence that auditory–motor learning relies on complementary processes operating over multiple timescales. By linking neural measures of prediction error to a skilled behavior, this work establishes a framework for studying how the brain continuously acquires, updates, and exploits internal models of sensorimotor interactions.

## Materials and method

### Participants

Twenty-one participants (12 female, ages 19-29, mean age 24.1, 3 left-handed) were recruited from the École Normale Supérieure community. Three participants were excluded due to technical anomalies during recording. Participants had various levels of musical training, ranging from no training to over 10 years of formal study. None of the musically trained participants played piano as their primary instrument. Written informed consent was obtained from all participants, and each participant was compensated for his/her participation. The study was conducted in accordance with the Declaration of Helsinki and was approved by the CNRIPH committee. Participants were briefed on the instructions by the experimenter and had access to a printed copy of the written instructions throughout the experiment.

### Experimental setup

Participants completed the experiment in a single session within a soundproof, electrically shielded booth with dim lighting. For all phases of the experiment, participants were instructed to fixate on a cross at the center of their visual field and minimize all motor activity except for the finger movements required to play the keyboard. A Novation Launchkey Mini MIDI keyboard was placed on the desk in front of the participant. Audio stimuli were presented monophonically at a sampling rate of 44.1 kHz using Sennheiser HD650 headphones at a volume comfortable for the participant. The experiment lasted approximately 3 hours, including 1 hour of setup, 1.5 hours of EEG recording, and short breaks between tasks at the participant’s discretion.

### Overview of tasks

Experimental sessions consisted of three main blocks (Figure S1), each lasting about 30 minutes: pre-training, training, and post-training. Pre- and post-training blocks were identical and consisted of three 10-minute tasks: *passive listening*, *mute playing*, and *variable-map playing*. Each of these tasks is detailed below.

### Variable map playing task

The participant’s right hand was positioned with the experimenter’s assistance on the ‘playing zone’ of the keyboard, comprising the keys F4, G4, A4, B4, and C5. One finger was positioned on each key. Textured fabric strips placed on the boundaries of the playing zone allowed participants to keep their fingers in the correct position throughout the experiment. Participants were asked to play 4-note sequences at approximately 60 beats per minute (bpm) separated by 2 seconds of pause, and to vary the sequences played.

The task lasted 10 minutes, during which the *key-pitch mapping* changed unpredictably every 2-10 seconds. Three mappings were possible as illustrated in Figure 2A: the *inverted* mapping (the playing zone keys, F4, G4, A4, B4, and C5, were mapped to G4, F4, E4, D4, and C4, respectively), *shifted-inverted* mapping in which pitches from the *inverted* mapping were shifted up one whole tone (to A4, G4, F4, E4, and D4, respectively), and *normal* mapping reflecting that of a normal piano (F4, G4, A4, B4, and C5). The complete assignment of mappings over the 10-minute playing session is detailed in Supplementary Figure S2. Sound synthesis and switching between mappings were automated using Ableton Live 11, a low-latency digital audio workstation for live music.

Participants performed the tasks with no visual input apart from the fixation cross to minimize eye movements. The beginning and end of each task were marked by a bell sound. The experimenter monitored the sequences played by the participant in real time from outside the experimental booth to ensure that participants followed the task instructions. The audio output was recorded and used in the control experiment described below.

### Training

With the right hand placed on the keyboard in the same way as in the variable map playing task, participants completed a self-guided training session (Figure 2) designed to familiarize them with one of the three key-pitch maps used in the variable map playing task. Throughout training, *key-pitch mapping* consistently followed the *inverted* mapping.

The training session consisted of 2 identical blocks, where the same 8 sets of 10-12 melodies were played (Figure 2D). The sets of melodies were arranged in ascending difficulty based on the number of pitches used in each melody (e.g., set 1 used only C4 and D4, while set 8 used all 5 keys). Each melody consisted of four notes, played at 60 bpm. Participants were asked to imitate the melodies played to the best of their ability. Melodies were isochronous and there was no penalty for rhythmic errors, but participants were asked to stay within the allotted response time of 5 seconds. There was an opportunity to take a break after each set. Played melodies were recorded using the MIDI protocol during both the pre- and post-phases, as well as during training. Training lasted approximately 30 minutes and included a total of 164 melodies to imitate.

Performance on the melody imitation task during training was scored by comparing the melody played with the target one, note by note, without considering timing. If participants answered with more than four key presses, only the first four were considered. If participants answered with fewer key presses, the response was padded with non-valid events. For each note, a score of 1 was awarded if the note was correct, and zero otherwise, meaning that the maximum score was 4 (Figure 2E). We observed a significant improvement in score for all participants (Figure 2F).

### Auxiliary tasks

Participants performed additional passive-listening and mute-playing tasks to generate subject-specific EEG data for training auditory and motor decoder models, as well as unimodal ERPs. The order of tasks was chosen to ensure that the listening and mute playing tasks during pre-training are not affected by participant expectations resulting from playing the keyboard (Figure 2C). Participants had the opportunity to take a break after each task.

In the passive listening task, participants listened to the corpus of melodies used in the training session. Each melody was presented once, with a 2-second pause between melodies, for a total of 11 minutes and 5 seconds of listening.

In the mute playing task, participants were asked to play 4-note sequences on a muted keyboard and to vary the sequence each time. The keyboard output (both audio and MIDI) was recorded in Ableton Live to ensure that participants played a variety of different melodies. The task lasted 10 minutes and was identical to the variable map playing task except that the participants heard no sound.

### Control experiment

To control for responses potentially related to auditory surprisal, six additional participants who did not participate in the original experiment were recruited to listen to the recordings of note sequences played during the full 10-minute variable map playing task. Eight recordings were used in the experiment, and each participant listened to a semi-random combination of 4 recordings such that each recording was heard by three participants. The sample size matched the number of note events analyzed in the original experiment while accounting for inter-participant variability in recordings and individual differences in neural responses. We did not invite the original participants to listen passively to their own playing to exclude the possibility that they recalled the played melodies.

### EEG acquisition and preprocessing

During the experiment, 64-channel EEG data, along with two external electrodes, were recorded at a sampling rate of 2048 Hz. The BioSemi ActiveTwo system and the accompanying software were used for EEG acquisition. In addition, 4 external electrodes (2 mastoid, 2 ocular) were used for re-referencing and eye-blink artifact removal, respectively. To ensure synchronization between EEG data and stimuli, all key presses on the MIDI keyboard, audio onsets, and trial start signals were routed to a customized analog trigger system, which was recorded as additional EEG channels.

EEG data were preprocessed and analyzed offline using MNE-Python (60) and following standard guidelines for linear modeling of neurophysiological data to auditory stimuli (61). EEG data were notch-filtered at 50 Hz and bandpass-filtered between 1 and 30 Hz using Butterworth zero-phase filters (order 3, forward and backward pass). All channels were re-referenced to the average of the two mastoid channels. Channels with a variance exceeding three times that of the surrounding ones were replaced by an estimate calculated using spherical spline interpolation (61). Finally, the data was downsampled to 128 Hz to reduce the computational load.

### ERP analysis

Data were segmented into ERP epochs using time-aligned note onsets (for motor-only tasks, the participant’s headphones were muted, but the corresponding audio information was still transmitted to the EEG acquisition computer). A pre-stimulus baseline (the average of activity between -500 ms to 0 ms) was subtracted from each channel in each epoch. Segments from 200 ms before to 500 ms after the note onset were isolated and averaged for analysis. Independent component analysis was performed separately on the 10 minutes of continuous data for each task (variable map playing, passive listening, and mute playing) and participant, rejecting components closely correlated with EOG signals. ERPs were classified as *firsts* or *others* based on their relative positions to map changes (Figure 2).

### Modeling of surprisal using IDyOM

To estimate the musical surprisal of individual notes, we applied the Information Dynamics of Music (IDyOM) model (20) using a Python implementation (31). The dataset consists of eight recordings, where each recording represents a participant’s variable map playing session in the main experiment. Surprisal values were computed for each note in each of the eight recordings using leave-one-out cross-validation: for each recording, the model was trained on the other seven recordings and then used to predict the note probabilities in the held-out recording. Both short-term (predicting probabilities based on up to 20 previous notes in the same recording) and long-term (trained on transition probabilities in all other recordings) models were used to calculate surprisal. Only pitch information was modeled; a uniform rhythmic structure was assumed, as participants did not vary the rhythm of the melodies.

### Statistical inference Permutation cluster tests

We first sought to identify time windows and channels in which ERPs differed significantly between first and other keystrokes. We performed a two-sided one-sample spatiotemporal permutation cluster test on the subject-level Δ*f − o*_ave_ difference waves (1,024 permutations), using a cluster-forming threshold based on the t-distribution. This non-parametric approach accounts for spatial and temporal dependence by identifying clusters of adjacent channels and time points. To assess whether the observed effects were also evident at the single-channel level, we performed a two-sided one-sample permutation cluster test on the difference wave at Fz, a channel previously implicated in auditory processing and ERPs associated with auditory onsets (9) (1,024 permutations). We further compared how the Δ*f − o*_ave_ difference waves were modulated by training, by computing Δ*f − o*_diff_ = Δ*f − o*_post_ *−* Δ*f − o*_post_. We then computed separate Δ*f − o*_diff_ waves for each key-to-pitch map, sorting keystrokes according to which map was active at the time of the keystroke. The same one-cluster permutation test was then performed on these difference waves. Finally, to assess the statistical significance of channel-wise differences between *first* and *other* ERPs, we performed Wilcoxon signed-rank tests on participant averages at each channel within the time window of interest (80–120 ms), with false discovery rate (FDR) correction across channels.

### Bootstrap comparison of *firsts* versus *others* keystrokes

To ensure that differences between *firsts* and *others* keystrokes found through 1-sample cluster permutation tests are robust to the class imbalance, *i.e.,* the fact that there are fewer *firsts* than *others* keystrokes, we employed a bootstrapping procedure at the time window of interest identified by the 1-sample permutation cluster test (0.08 to 0.12 s). For each of the *N* = 1000 bootstrap iterations, we drew *n* = 100 random samples with replacement from the set of *firsts*, and *n* = 100 from the set of *others* for each subject. We then computed the mean ERP amplitude for each class at the time window and calculated the within-subject difference between the two. The average difference across subjects was retained at each iteration, yielding a bootstrap distribution of mean differences across subjects. To generate a null distribution, the same procedure was applied using two random samples of *n* = 100 trials each drawn from the *others* keystroke class, thus controlling for differences unrelated to keystroke category. Statistical inference was based on comparing the empirical and null bootstrap distributions: we evaluated whether their 95% confidence intervals (CI95) overlapped, and whether either distribution’s CI included zero. We applied the same bootstrapping procedure to Fz, Cz, and Iz electrodes to confirm that the observed distributions of mean differences were consistent with the spatial patterns seen in the ERP difference topographies, deliberately including two channels in the region of interest as identified by previous auditory-motor literature (9) (Fz, Cz) and one channel well outside the region of interest (Iz).

### Keystrokes contextual analysis

To assess whether auditory-motor surprise is modulated by the previous context, we analyzed N100 amplitudes of *first* keystrokes as a function of the number of keystrokes performed in the map preceding the map change. We defined a *0-keystroke* context as the case where the note immediately preceding the current first keystroke was itself a first keystroke. A *1-keystroke* context corresponds to a case in which one *other* keystroke (i.e., a keystroke *not* following a map change) precedes the first keystroke and a first keystroke occurs two keystrokes prior, and so on. Trials were pooled across participants, and only classes of first keystrokes with more than 300 samples were retained to ensure a sufficient signal-to-noise ratio. Due to frequent map changes, longer context sequences were increasingly rare and thus excluded. This threshold on the number of samples allowed us to analyze up to a context of 3 keystrokes. To determine whether *first* keystrokes elicited larger N100 amplitudes when the *first* comes after a large number of *other* keystrokes before the map change, we fit a linear mixed-effects model on N100 amplitude of *first* keystrokes, with keystroke context as a fixed effect and subject as a random intercept. Using these models allowed us to leverage the large number of keystroke-level observations while accounting for between-subject variability. Relatedly, to determine whether the number of intervening keystrokes since the first keystroke of the map has an effect on N100 amplitude of *other* keystrokes, we fit a linear mixed-effects model for N100 amplitude of *other* keystrokes, with preceding keystrokes as a fixed effect and subject as a random intercept. Again, only keystroke classes with more than 300 samples were retained, allowing us to analyze up to 7 intervening keystrokes.

### Decoding analysis

Subject-specific decoders were trained separately using EEG responses to the passive listening and mute playing tasks (time lags -0.5 to 0.5 s). The *auditory decoder* was trained on EEG responses to passive listening to reconstruct note onsets, while the *motor decoder* was trained on EEG responses to mute playing to reconstruct keystrokes. Both decoders were then used to predict note onsets and keystrokes from EEG responses to the variable map playing task (Figure 7). Separate auditory and motor models were trained and tested for each participant using data exclusively from that individual. Amplitudes of the reconstructed onset vector were extracted at the ground-truth times of *first* or *other* keystrokes, and averaged separately to produce a mean predicted amplitude for *firsts*, and for *others*.

## Data and software

Statistical analyses were performed in Python using scipy (62), pingouin (63), and statsmodels (64). Significance stars in figures were automatically drawn using statannotations (65). Decoders were trained using mTRFpy (66).

## Data availability

The code for analyzing data and generating the figures is available on Github at https://github.com/haiqin-zhang/AuditoryMotorPiano. EEG data is available on Zenodo (DOI: 10.5281/zenodo.17949131).

## Acknowledgments

This work was supported by an Advanced European Research Council grant (NEUME, 787836), FrontCog grant ANR-17-EURE-0017, and AFOSR grant (FA9550-24-1-0242) to SS. HZ is supported by a *Contrat Doctoral Spécifique Normalien* (CDSN) doctoral grant funded by the French Ministry of National Education, Higher Education, Research and Innovation, and administered by École Normale Supérieure – PSL. The project was based on pilot experiments and discussions at the 2023 Telluride Neuromorphic Engineering Workshop. The authors are grateful to Yves Boubenec, Giovanni Di Liberto, Claire Pelofi, and Virginie van Wassenhove for their feedback during conception and writing.

## Supplementary figures

**Figure S1:**
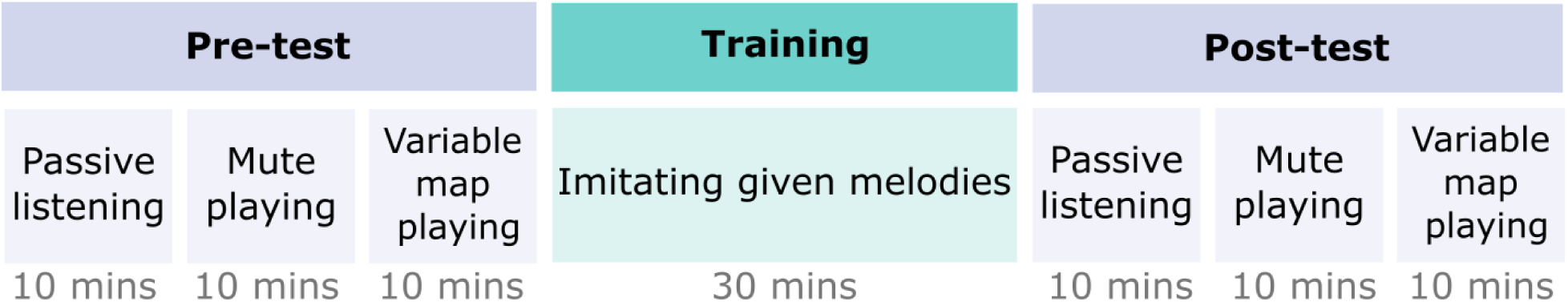
Overall experimental procedure showing order of the variable map playing, listening, and mute playing tasks carried out both before and after the training task.

**Figure S2:**
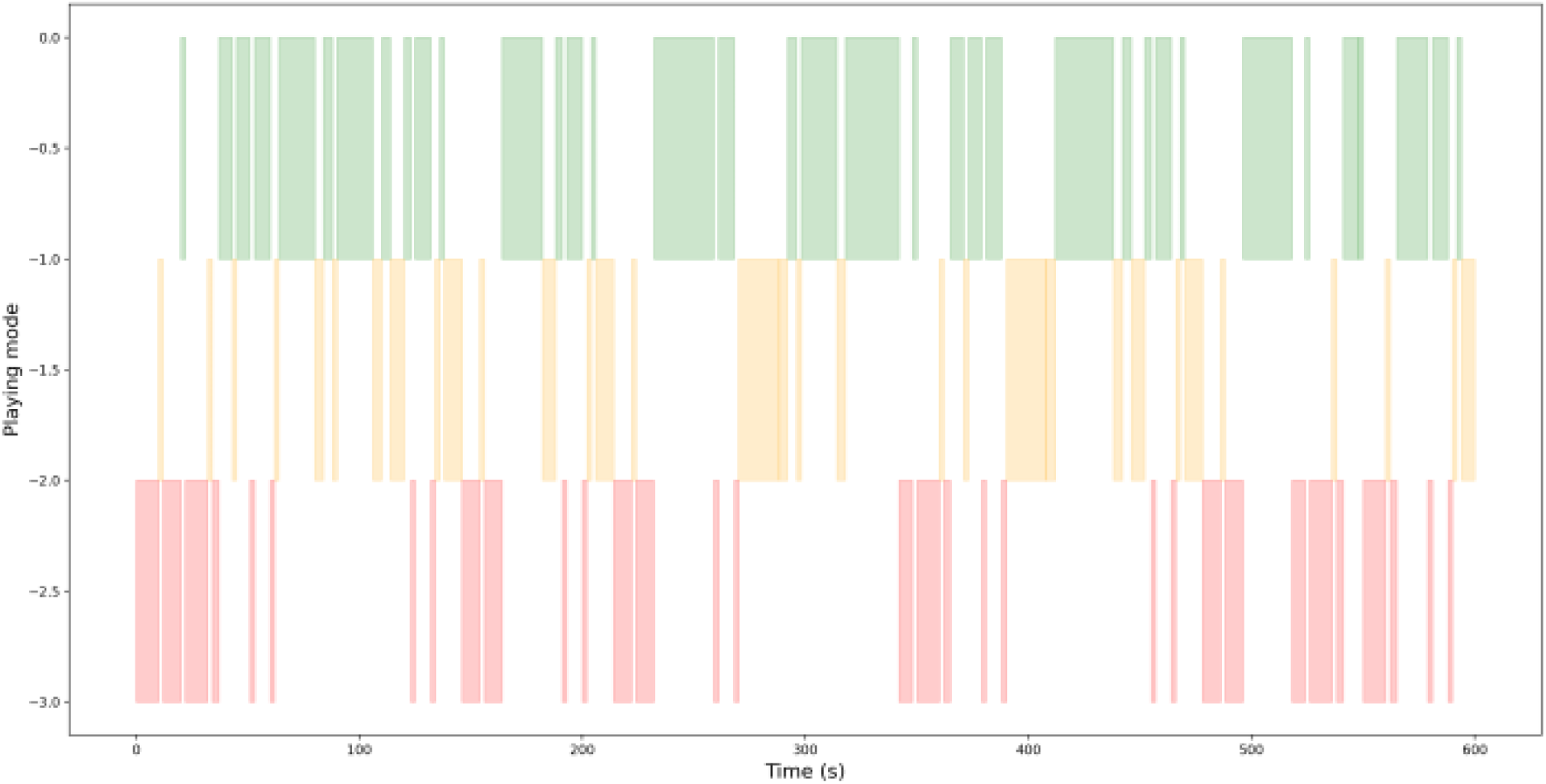
Key-pitch map assignment over time during the full 10-minute task.

**Figure S4:**
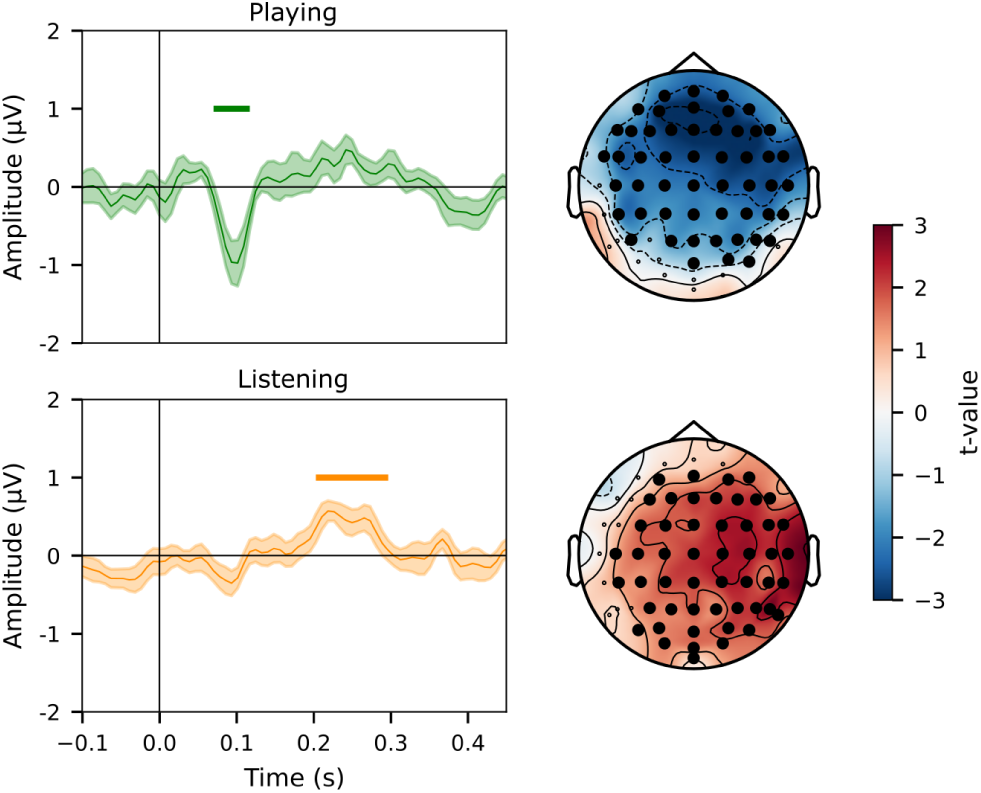
Grand average note onset ERPs during the passive listening, mute playing, and variable map playing tasks in all channels, in pre-training only. Colours represent EEG channels.

**Figure S3:**
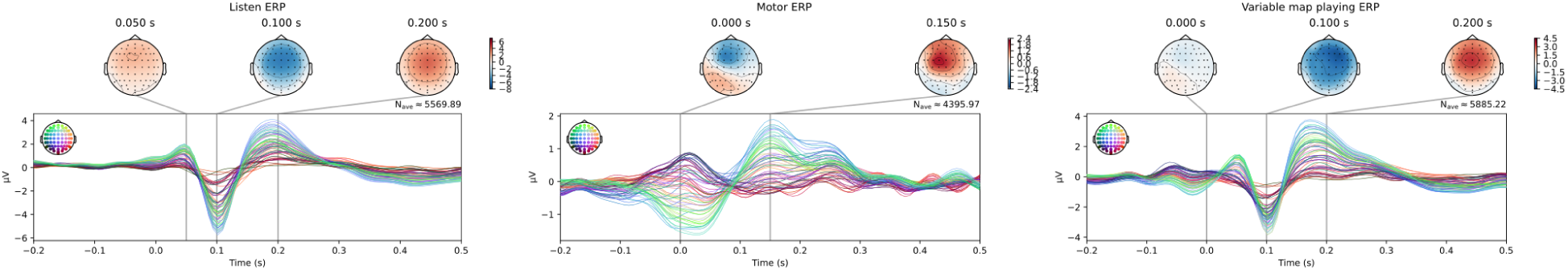
Significant spatiotemporal clusters for first vs. other keystrokes during playing and listening. Lines show ERPs averaged across significant channels (bold in topomap); shaded areas indicate SEM. Bars indicate significant temporal clusters. *p <* 0.05, one-dimensional spatiotemporal permutation cluster test.

**Figure S5:**
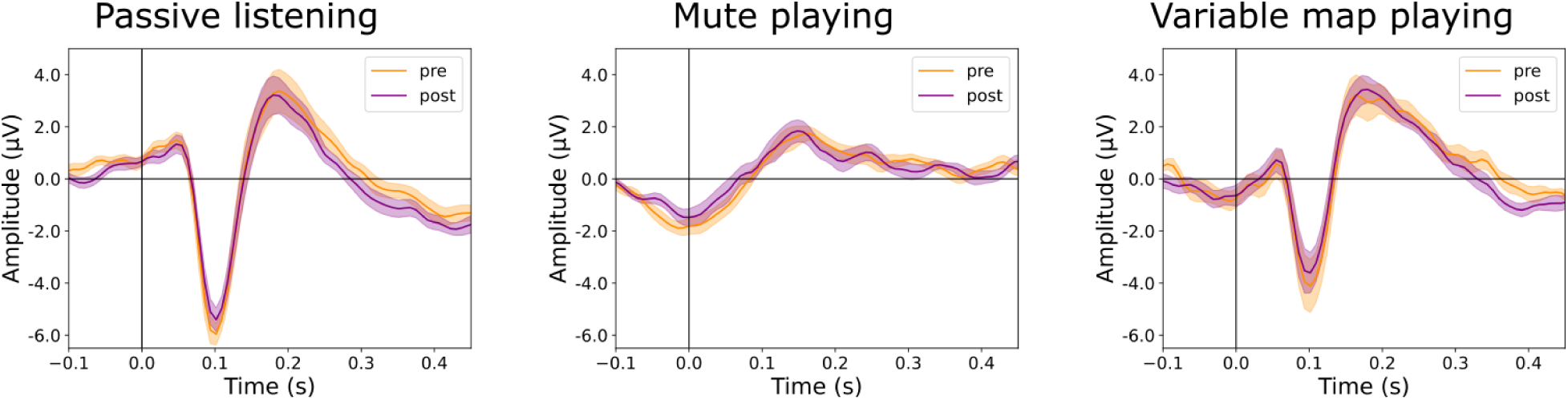
ERPs aligned to the note onsets during the passive listening, mute playing, and variable map playing tasks in the FCz channel, showing differences pre and post-training. Lines represent the grand average over all subjects, shading represents SEM.

**Figure S6:**
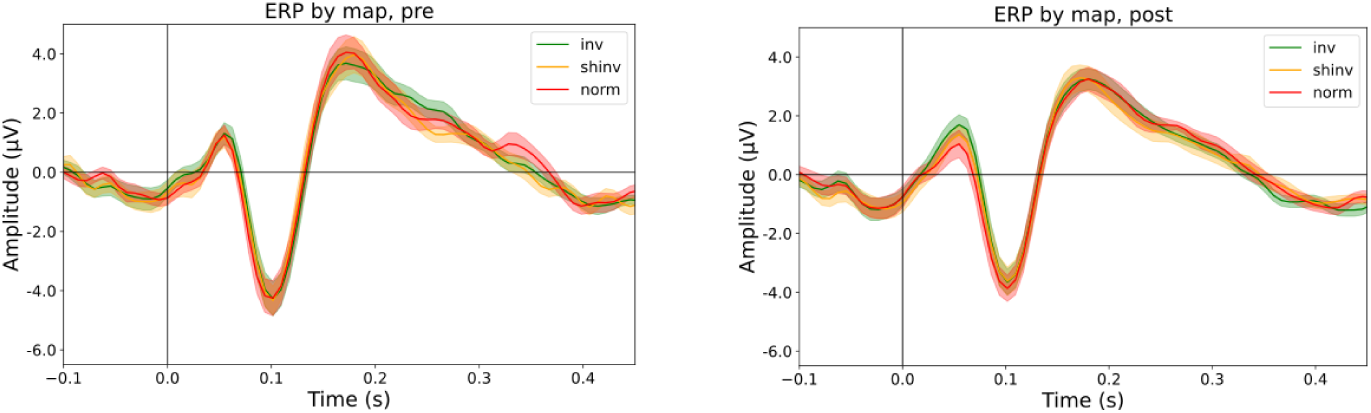
Note onset ERPs sorted by map, including all *firsts* and others, separated by pre and post-training. Lines represent the grand average over all subjects, shading represents SEM.

**Figure S7:**
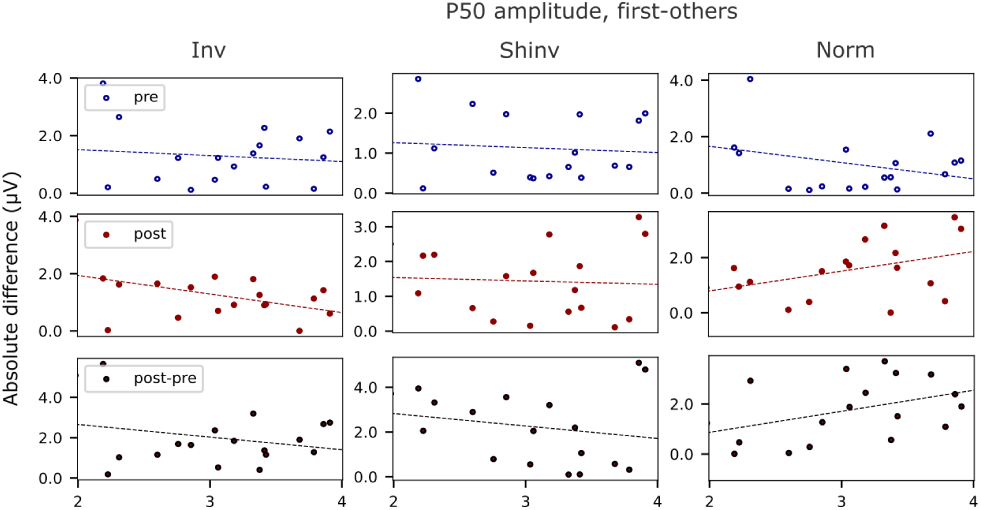
Correlation between training score and Δ*_f−o_* at P50. Columns show the active key-pitch map at the time of the keystroke. Rows show correlations in pre-training, post-training, and the difference between the two (post-pre). Dotted lines show linear regression. *p >* 0.05 for all panels, Pearson’s correlation.

**Figure S8:**
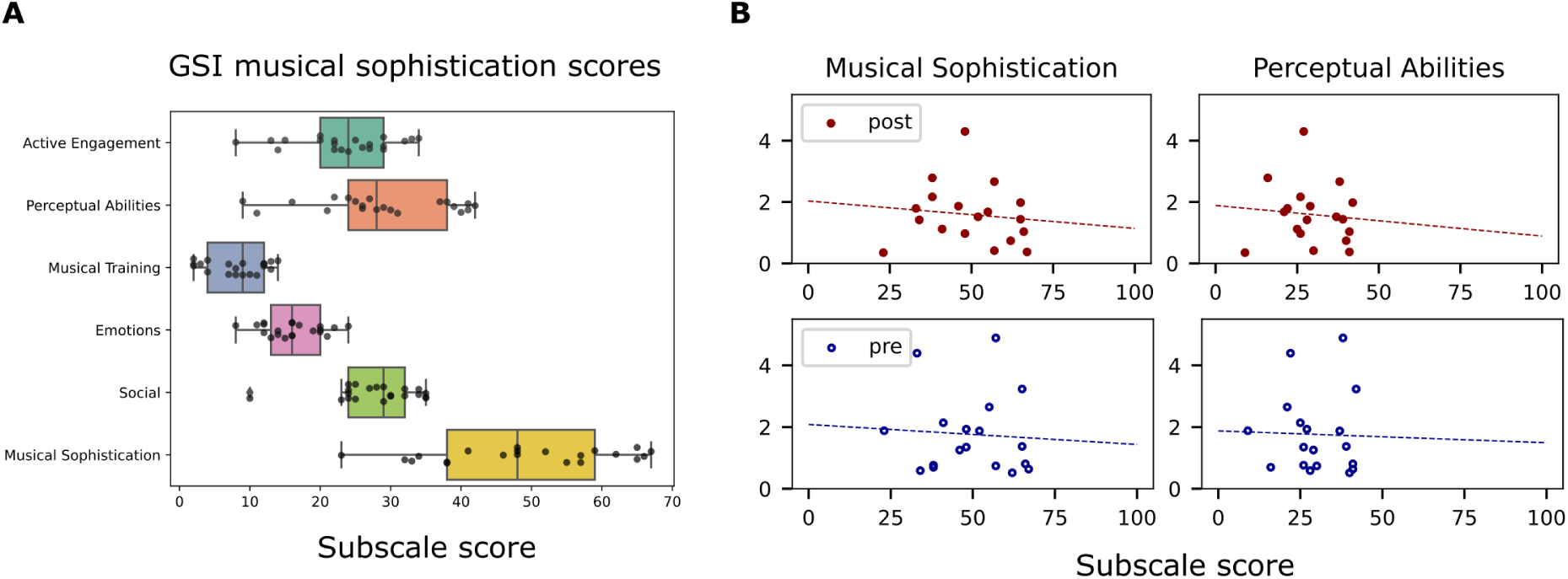
Goldsmiths Muscial Sophistication Index scores and correlation Δ*_f−o_*. **A.** Distribution of scores across subjects on overall musical sophistication, and sub-scale categories. Boxes represent median and quartiles, points represent individual subjects. **B.**Correlation between Δ*_f−o_* at P100 and overall musical sophistication score (left), or perceptual abilities sub-scale score. *p >* 0.05 for all panels, Pearson’s correlation.

**Figure S9:**
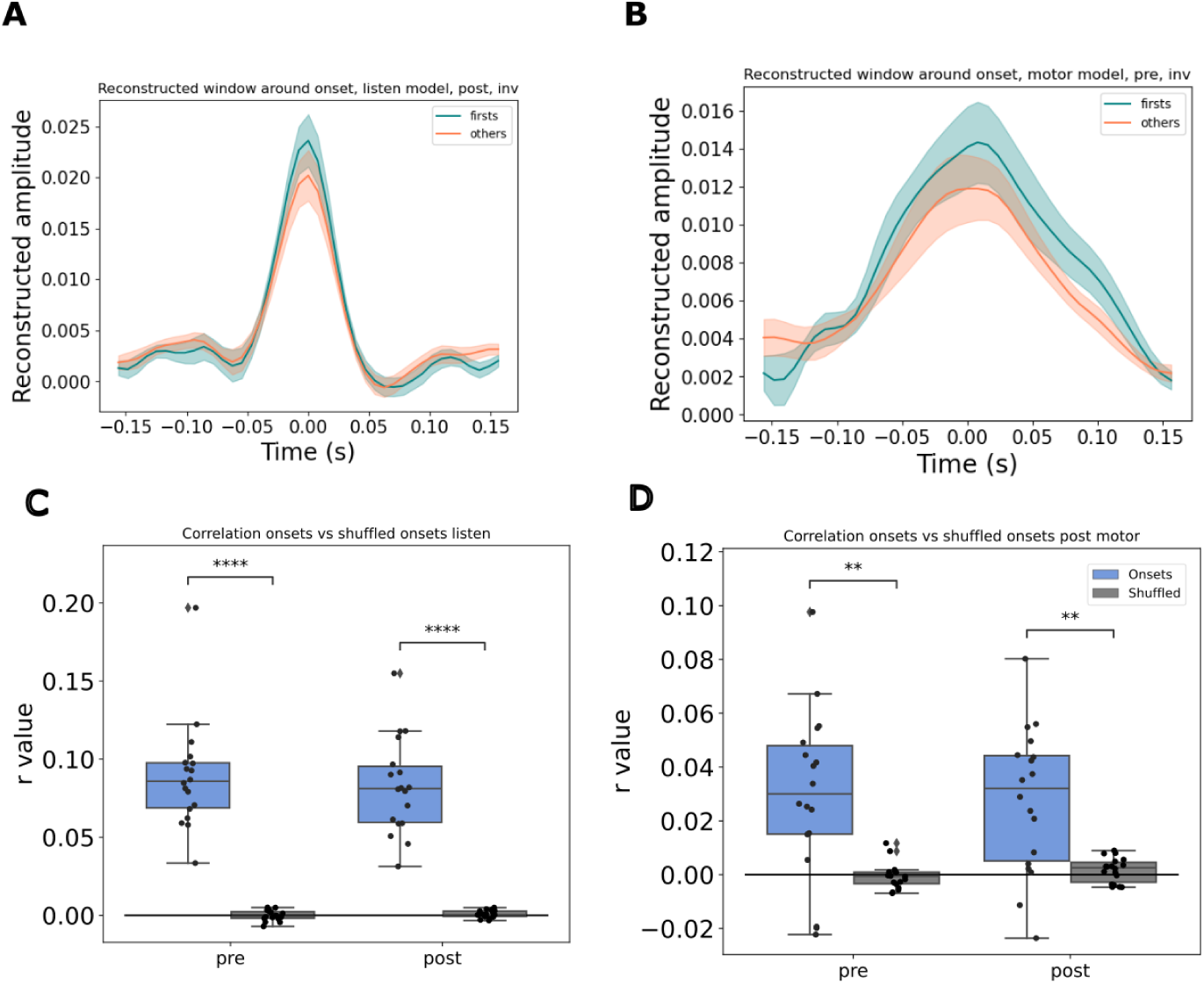
Reconstruction accuracy of inverse TRF decoder. **A-B.** Average window around note onset times in reconstructed stimulus using listening and motor decoders, respectively. **C-D.** Correlation between ground truth (sparse vector with time of note onsets set to 1) and reconstructed stimulus when the ground truth is shuffled versus unshuffled, for listening and motor decoders, respectively. ** *p <* 0.01, **** *p <* 0.0001, Pearson’s correlation.

**Figure S10:**
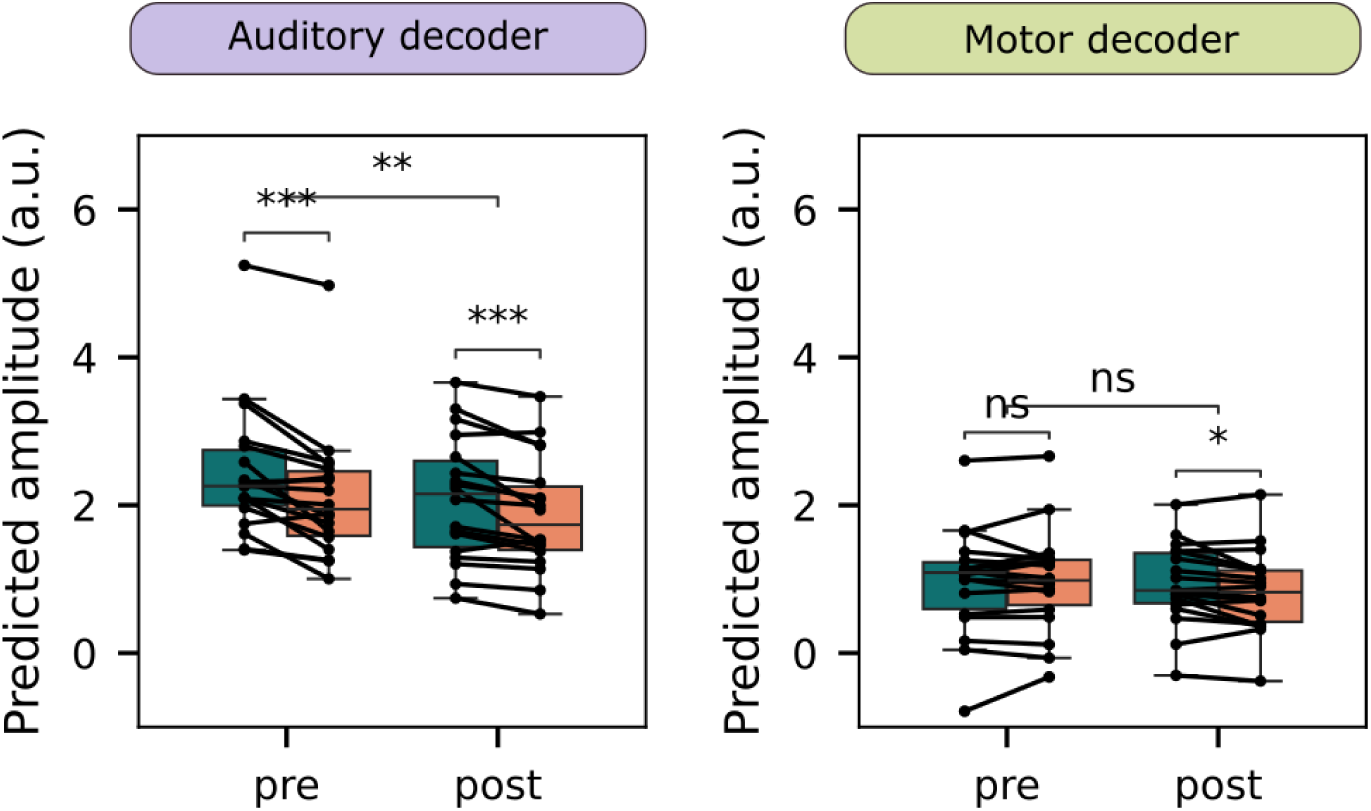
Reconstruction amplitudes at first and other keystrokes, with averages computed from size-matched samples. Lines represent participants; bars show medians and quartiles based on 100 samples of firsts and others per participant. Wilcoxon signed-rank test, p < 0.001.

**Figure S11:**
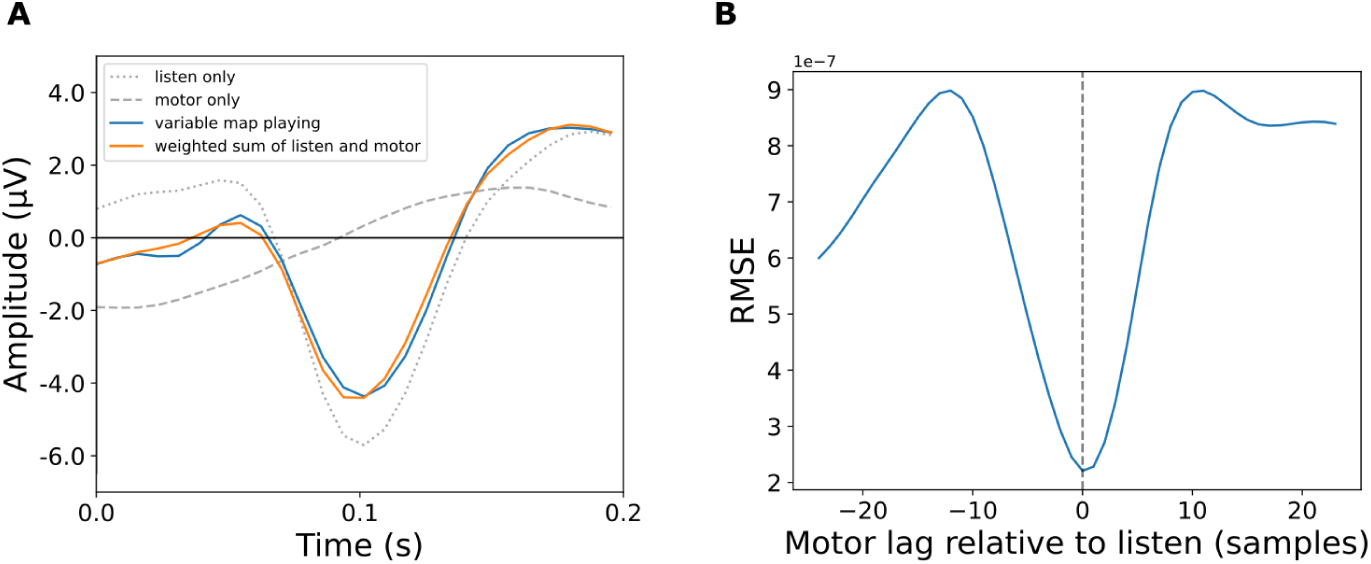
Comparison of weighted sum of auditory and motor ERPs with playing ERP. **A.** Grand average ERPs for the auditory-only condition (locked to note onset), motor-only condition (locked to keystrokes), the variable map playing ERP, and the optimized weighted sum of auditory and motor ERPs. Weights were determined by least squares optimization. **B.** Root mean square error (RMSE) between the playing ERP and the optimized weighted sum of auditory and motor ERPs across different lags applied to the motor ERP relative to the auditory ERP. The lowest RMSE occurs at zero lag.

**Supplementary video.** Excerpt of a variable map playing session. Audio is played by the participant. Each row represents one key-pitch map (green: *inverted*, yellow: *shifted-inverted*, red: *normal*). Bars in that row represent the beginning of that map (and the end of the previous one).

## Footnotes

1 For clarity, we shall refer to the *external* movement-to-sound correspondence as the *key-pitch* map, and its neural representation as the *auditory-motor* map. The key-pitch map is determined by external factors (e.g., the piano configuration), while the auditory-motor map exists only within the brain’s sensorimotor system and is adjusted through learning.

